# Extracellular Thimet Oligopeptidase is Released with Extracellular Vesicles from Human Prostate Cancer Cells

**DOI:** 10.1101/2020.05.10.087304

**Authors:** Yu Liu, Lisa Bruce, Adele J. Wolfson

**Author notes:** Correspondence to: Adele J Wolfson, Wellesley College, Department of Chemistry, Wellesley, MA 02481.

## Abstract

Androgen signaling plays a central role in the development of prostate cancer. Androgen hormone synthesis is tightly governed by the hypothalamic–pituitary–gonadal (HPG) axis, including gonadotropin-releasing hormone (GnRH). Thimet oligopeptidase (TOP) is a biologically significant peptidase known to cleave GnRH and potentially regulate its activity. Thus, TOP can play an important role in the HPG axis through regulating the downstream production and release of gonadal steroid hormones, including androgens, which may further affect prostate cancer development. TOP is known to be secreted out to the extracellular space. Here, we report that extracellular TOP can be associated with extracellular vesicles (EVs). Western blot analysis of EVs isolated from PC3 or DU145 prostate cancer cells revealed that TOP protein is, indeed, carried by the EVs. Budding of EVs from stimulated PC3 prostate cancer cells can also be visualized by confocal microscopy. Significantly, the TOP enzyme carried by EVs is enzymatically active. The present study shows that EV-associated TOP is a novel form of this extracellular peptidase that may play a role in the disease progression of prostate cancer cells.

## 1. Introduction

Prostate cancer is the second leading cause of cancer-related death in men in the US. It is estimated that 1 in 6 men will be diagnosed with prostate cancer and approximately 1 in 33 (3%) will die of it^1^. It has long been known that androgen signaling plays a central role in the development of prostate cancer^2, 3^ and androgen deprivation therapy (ADT) remains a major therapeutic strategy for advanced prostate cancer^2–4^. Androgen hormone synthesis is tightly governed by the hypothalamic–pituitary–gonadal (HPG) axis. The pulsatile release of gonadotropin-releasing hormone (GnRH) stimulates the secretion of luteinizing hormone (LH) from the anterior pituitary gland, which then signals the testes to produce testosterone, the major androgen hormone in men. Therefore, GnRH is the major regulator in androgen production^3^. GnRH antagonists act as competitive inhibitors of GnRH and decrease the secretion of LH from the pituitary, thereby leading to a decrease in testosterone secretion from Leydig cells in the testes^5^. Furthermore, GnRH antagonists are currently FDA-approved frontline treatments of prostate cancer and are widely used in clinical applications^39^.

Thimet oligopeptidase (TOP, E.C.3.4.24.15) is a well-characterized zinc metallo-endopeptidase that hydrolyzes only short peptides with less than 20 residues, and recognizes greatly variable cleavage sequences depending on the specific substrate^6–9^. Studies have indicated that GnRH is a substrate of TOP^6, 10^, which can cleave and regulate activity of GnRH^9–11^. Via regulation of GnRH, TOP modulates the hypothalamus–pituitary–gonadal axis as well as the downstream production and release of gonadal steroid hormones, like testosterone^11, 12, 41^. Furthermore, TOP is highly expressed and regulated in prostate tissue, and its actions are increased in androgen-dependent prostate cancer compared to androgen-independent cancer^11^. Thus, TOP is involved in both etiology and pathophysiology of prostate cancer^11^.

TOP has been found to be distributed in different subcellular locations depending on cell types, i.e. nuclear localization^13, 14^, membrane association^15–17^, cytosolic and secreted forms^13–16, 18–21^. Most of peptide processing is initiated in the intracellular space of parental cells,^6^ while many peptides need to be further processed and activated in the extracellular milieu by enzymes, such as TOP^6^. Although cytosolic TOP account for 75% of enzyme activity, a significant portion of TOP released to the extracellular milieu is responsible for the extracellular processing of neuropeptides such as GnRH^22^. In fact, studies show that TOP can be secreted to the extracellular surroundings^6, 18, 19^ and TOP activity has been detected in the media of DHT-treated LNCaP androgen responsive cells^11^.

Extracellular vesicles (EVs) are small cellular membrane vesicles that are either released from intracellular space (exosomes)^27^, or budded from the plasma membrane surface of almost all cell types upon cell activation or programmed cell death (microvesicles)^23–27^ In the past several years, the biomedical research field has shown growing interest in the biological roles of EVs due to their emerging roles in pathophysiological conditions and various human diseases^26, 27^Studies show that EVs carry a heterogeneous array of intracellular and membrane-associated biologically active molecules, such as proteins, lipids, and DNA or RNA^23–27^. Previous studies demonstrate that EVs carry transmembrane protease of the matrix metalloproteinase superfamily (MMP14)^23^. Given that TOP has been visualized on the cell plasma membrane surface by confocal microscopy^17^, we sought to investigate if TOP can be released with EVs, in addition to its free form^11^, to the extracellular space. Soluble molecules can be diluted by the large pool of circulating blood, while EV-associated molecules can be kept in a relatively concentrated microenvironment around the parental cells that release these EVs. Additionally, EVs can interact with cells/tissues across vast distances^28, 29^. Therefore, we propose that EV-associated TOP may be important in GnRH processing and activity regulation in a systemic manner.

## 2. Material and methods

### 2.1. Cell culture and treatment

The PC3 or DU145 cell lines were cultured at 37 °C and at 5% CO2 and grown in RPMI-1640 medium with 10% Fetal Bovine Serum at pH 7.8. PC3 cells were treated without (control) or with 5, 10, or 20 μM calcium ionophore (A23187) for 24 hours. DU145 cells were treated with 0.3 μM DHT or its vehicle (ethyl alcohol) for 24 hours. Cell lysates were collected to extract TOP, while the supernatant was harvested for EV purification.

### 2.2 Purification of MVs from cell culture supernatants

The supernatants from above control and treated cells were initially centrifuged twice at 3000 rpm for 15 min to remove residual cells, following by ultracentrifugation at 100,000 g for 30 minutes (Gyorgy et al., 2011; Li et al., 2011, 2012). Then the EV pellets were re-suspended in PBS with 2% SDS.

### 2.3 Western blotting analysis of TOP protein

Western blot was conducted, as described before^23^, for EVs (at the maximum lane volume) to detect whether the EVs carry TOP protein, cell lysates (10 μg/lane) were used as control. Transferred membrane was blocked in 5% milk, followed by overnight immuno-probing at 4°C with 1:300,000 anti-TOP (rabbit IgG) and 1:250,000 anti-actin (mouse IgG) primary antibodies. Secondary anti-TOP and anti-actin antibodies (GE Healthcare) were used at 1:10,000. The probed membrane was visualized using the Storm Scanner apparatus. Band intensity was quantified using ImageQuant software.

### 2.4 Immunocytochemistry analysis

Cells without or with A23187 treatment were blocked in 20% normal goat serum (NGS) and 1% BSA in 0.05M TBS, then incubated in 1:5000 monoclonal Rabbit-anti-TOP antibody with 1% NGS in 0.05M TBS with 0.02% sodium azide, 0.1% gelatin (pH 7.6) for 24 hrs. The cells were washed with 0.05M TBS with 0.02% sodium azide, 0.1% gelatin, and 10% Triton-X, then incubated in secondary Donkey-anti-Rabbit Red Alexa 594 (Invitrogen) at 1:100 and FITC-labeled Annexin V with 1.5% NGS in 0.05M TBS with 0.02% sodium azide and 0.1% gelatin for 90 minutes, then washed with the same buffer, and staining with DAPI for 30 minutes. The slides were cover slipped with Gel/Mount (biomeda, cat# M01). Imaging: The cells were imaged using a Leica TCS SP5 II confocal microscope.

### 2.5 Assessment of TOP Activity with Quenched Fluorescence Assay

Cell lysates and purified EV suspensions were used for TOP activity assessment. The assay buffer was 25mM TRIS HCl (0.125mM KCl, 1μM ZnCl2, 1mM TCEP, 10% glycerol). Wild type TOP (0.1μM) was used as assessment functioning control. TOP substrate Methyl Coumarinyl 18 Acetate (MCA) was used at a concentration of 1.4 mM. The competitive inhibitor of TOP, N-[1-(R,S)-carboxy-3-phenylpropyl]-Ala-Ala-Tyr-p-aminobenzoate (cFP) was used at a concentration of 16.5 μM to assess breakdown in prostate cancer cells not due to TOP cleavage. All fluorescence assays were conducted using a Molecular Devices SpectraMax M3 microplate reader with emission wavelength at 400nm, and excitation wavelength at 325nm. Activity was monitored every minute for up to 30 minutes. The slope of the graph (fluorescence intensity/minute) was used to determine the activity of TOP. The activity from samples was subtracted from the blank, which consisted of 1ul of MCA and 199ul of assay buffer.

## 3. Results

To address whether extracellular TOP is associated with EVs, cell models were established by treatment of cells using calcium ionophore A23187 which is known to induce EV release from cells by increasing intracellular calcium and membrane phospholipid redistribution^30,31^. Therefore, PC3 cells were stimulated with 5 or 20 μM A23187 to induce cell activation and EV release (Fig. 1A). The culture supernatant samples from untreated control and A23187-treated cells were subjected to centrifugation twice with low-speed to remove any floating cells and cell debris, followed by ultracentrifugation to isolate EVs from supernatant of two treatment groups and the control group. Western blot analysis showed TOP being detected within the isolated EVs and their parental cell lysates of both A23187-treated and control cells (Fig. 1A). Results indicate that the amount of TOP were undetectable on EVs from cells that were treated with 5 μM A23187 and those of the control group. In contrast, the EVs from cells that were treated with 20 μM A23187 showed a clear TOP protein band (Fig. 1A) as compared to corresponding TOP bands from their parental cells and the positive control recombinant TOP protein (Fig. 1A). In support of the above findings, we found that stimulation of PC3 prostate cancer cells with 20 μM A23187 can cause up to 90% of cells to undergo apoptosis (not shown). This is consistent with the observed phenomenon that EVs may be released by cells undergoing apoptosis. Furthermore, this is in line with previous studies showing that tobacco smoke extract treatment can cause human macrophages to undergo apoptosis and release EVs that carry matrix metalloproteinase-14 protein^23^.

**Figure 1.**
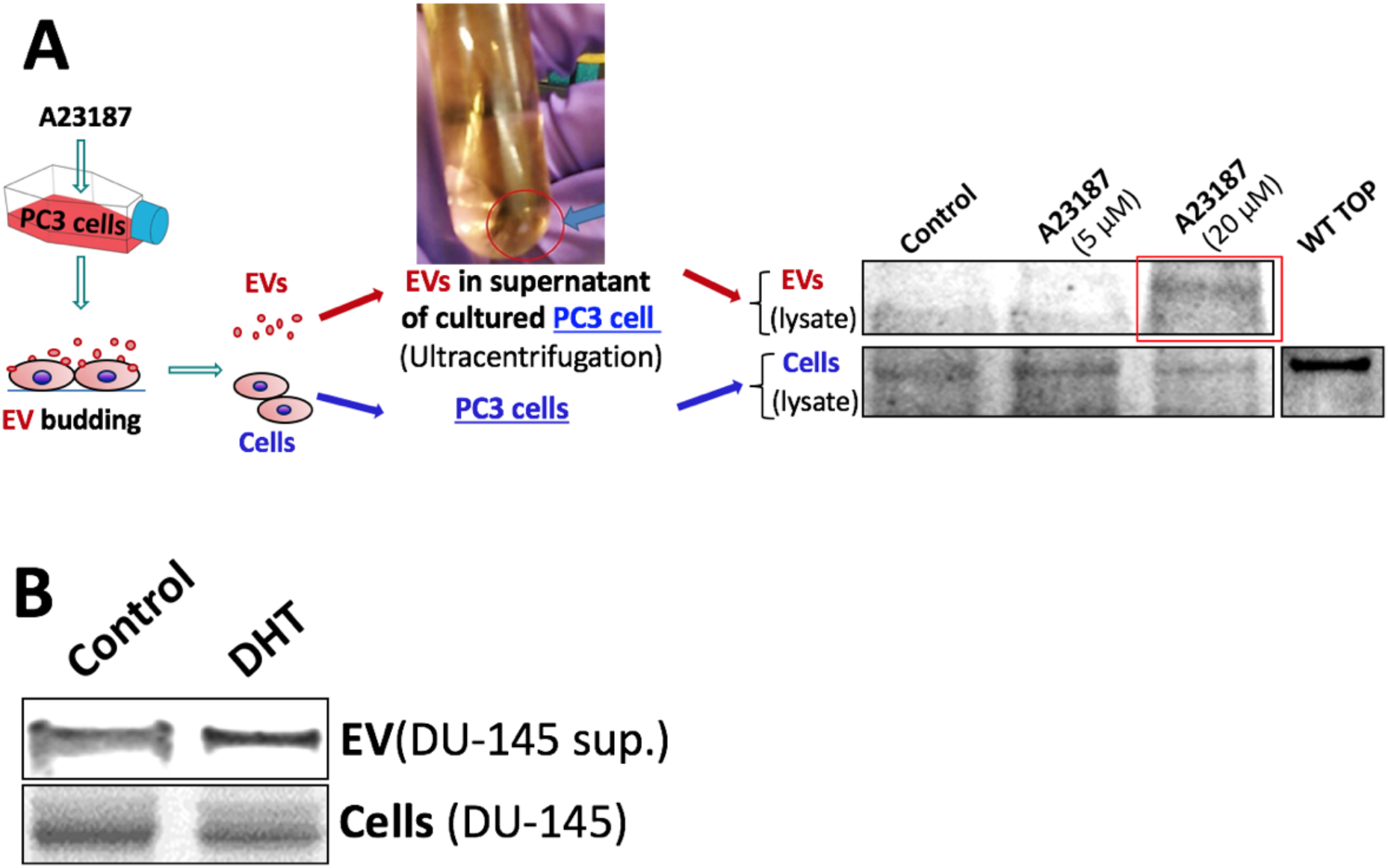
Detection of TOP molecules in isolated EVs from PC3 and DU-145 cells. **(A)** The PC3 cells were treated without (untreated control) or with 5 or 20 μM A23187 for 24h, then the EVs isolated from supernatants (sup.) or from the cells were lysed and analyzed via Western blotting. Recombinant wild type (WT) TOP served as positive control for detection of TOP. (**B**) The DU145 cells were either untreated (control) and treated with 0.3 μM DHT for 24h, then the EVs isolated from supernatants (sup.) or the cells were lysed and analyzed via Western blotting. TOP was probed by rabbit anti-human TOP primary antibody.

Having detected TOP expression in DU-145 cells, another prostate cancer cell line^32, 33^, we next sought to determine whether the EVs released from these cells also carry TOP protein. Immunoblot results indicated that EVs from the culture supernatants of both control and the DU-145 cells treated by 0.3 μM DHT do indeed carry TOP protein (Fig. 1B). Therefore, the fact that TOP enzyme is carried by EVs in the extracellular space helped us to confirm our initial hypothesis that extracellular TOP can be associated with EVs, and not only in the soluble form. However, this novel finding is not surprising since EVs bleb off of the plasma membrane at random locations, where TOP molecules may be associated with the membrane surface.

After detecting TOP molecules from the isolated EVs, TOP in both EVs and cell membrane surface was further visualized by confocal microscopy of PC3 cells that were treated without (control) or with 10 μM A23187 for 24 hours. Interestingly, the confocal cross-sectional images highlighting the cellular surface of PC3 cells not only showed membrane-associated TOP, but revealed co-staining of phosphotidylserine-rich domains in small, well-circumscribed blebs (Fig. 2A). Phosphotidylserine is a marker of EV generation^34^. The sizes of these membrane surface domains are consistent with the recognized size range for membrane microvesicles (approximately 0.1 to 1 μm)^26,27,35^. Thus, A23187 treatment appears to induce the budding and release of EV-associated TOP protein. Furthermore, the images show a concentration of cellsurface TOP induced on the cell surface of A23187-treated cells compared to the untreated control cells (Fig. 2A). These results confirm and extend our findings with western analysis of the isolated EVs (Fig. 1A-B). EVs are known for their heterogeneity and carriage of various molecular cargos. This heterogeneous nature is clearly demonstrated in Figure 2A, which shows a large concentration of membrane-surface TOP as a result of induction apoptosis by A23187. The visualized, circumscribed cell-surface domains that stained intensely for both TOP and externalized phosphotidylserine also suggest that the TOP-associated EVs are budding from apoptotic cell surfaces (Fig. 2A). Furthermore, as seen in the merged image, co-localization of TOP and phosphotidylserine is variable, thus illustrating the heterogeneity of EVs and the array of molecular cargo they carry.

**Figure 2.**
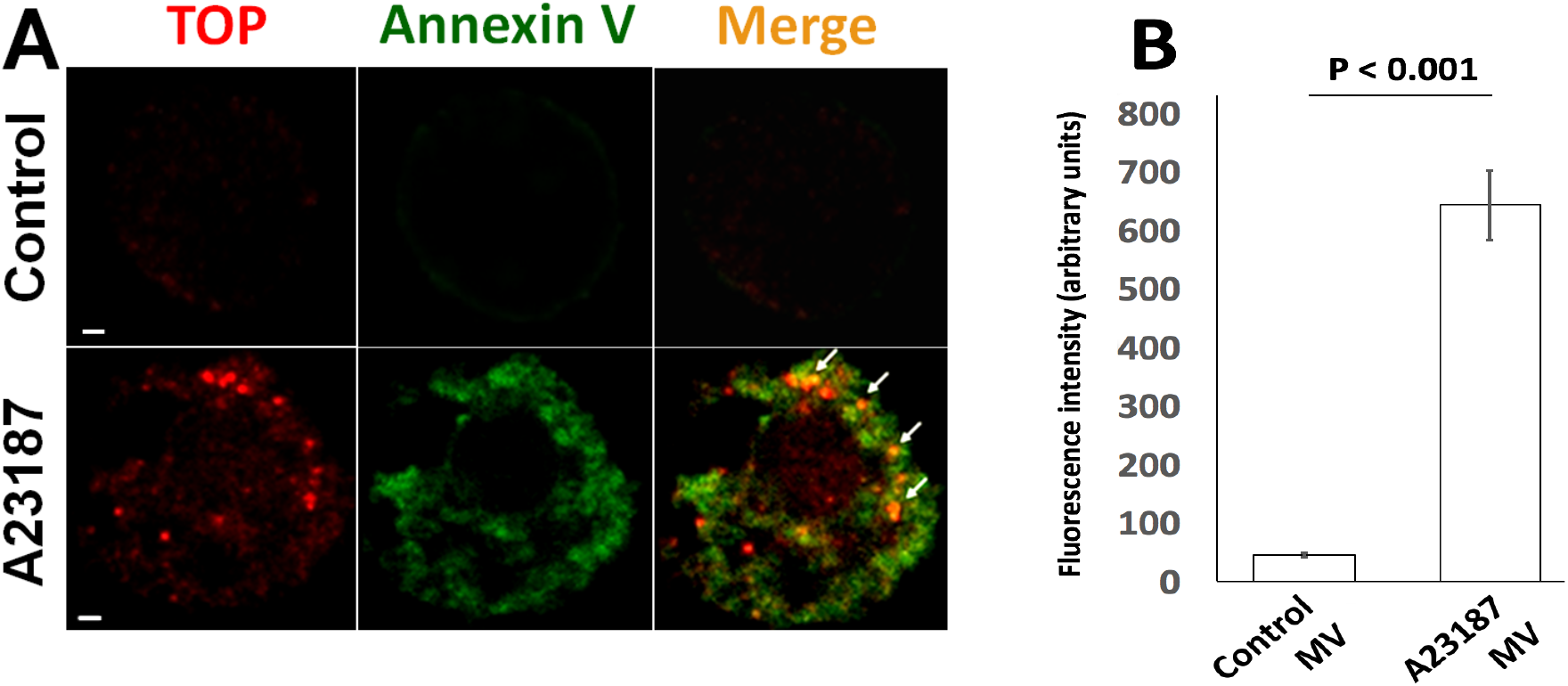
TOP-positive EVs were budding from plasma membrane surface of A23187-activated PC3 prostate cancer cells. **(A)** Cells were either untreated (control) or treated with 10 μM A23187 for 24 hrs. Cells were then chemically fixed and detected with anti-TOP antibody (**Red**, Alexa 594 labeled) and FITC-labeled Annexin V (**Green**, which preferentially binds externalized phosphatidylserine). The **yellow** color in the merged images (Merge) demonstrates co-localization of the two labels indicating the TOP-positive EVs. Scale bar 1 μm. (**B**) To assess TOP activity in cells and the isolated EVs, cells were treated without (control) or with 10 μM A23187 for 24 hrs. Then the cell lysate and EV suspension were used for detection of TOP activity. Enzymatic activity of TOP was determined using quenched fluorescence kinetic assay, by the cleaving of fluorescent-tagged artificial substrate (7-methoxycoumarin-4-yl) acetate (MCA). Enzyme rates, as depicted above, were obtained by calculating the slopes (fluorescence intensity, arbitrary units). Error bars were calculated using standard deviation.

Finally, we sought to evaluate whether TOP carried by EVs is enzymatically active. The purified EV samples isolated from A23187-treated or control cells were then analyzed via TOP enzymatic assay. Enzyme rates were calculated in terms of change in fluorescence per unit time. Robust TOP activity was detected in the EVs isolated from A23187-treated prostate cancer cells as compared to the EVs from the untreated control cells (Fig. 2B). These interesting findings not only confirm the presence of TOP protein on membrane EVs, but also clearly demonstrate that the EV-associated form of extracellular TOP is enzymatically active.

## 4. Discussion

TOP is generally considered to be a soluble molecule in the extracellular milieu^11^. However, the current study highlights for the first time a previously unrecognized novel form of extracellular TOP that is carried by the membrane EVs. Most importantly, we showed that extracellular EV-associated TOP exhibits considerable enzymatic activity (Fig. 2B). We have shown that EV-carried extracellular TOP can be released from two different human prostate cancer cell lines, both PC3 and DU145 prostate cancer cells (Fig. 1A-B). The DU145 results suggest that EV-associated TOP may also be released under natural conditions without additional stimulation with DHT hormone (Fig. 1B). As noted, many physiological substrates of TOP, such as GnRH, bradykinin and neurotensin, are found either in the extracellular space or in the systemic circulation. In order to fully modulate its physiological substrates, TOP must be accessible to the extracellular space, either by exposure on the cell membrane surface or via release from the cell as a soluble protein. Previous studies have localized TOP to the cell surface of the plasma membrane^17^ and have shown that TOP is secreted to the extracellular surroundings as a soluble free protein^6,18,19^. The current study highlights a previously unrecognized, novel form of extracellular TOP that is carried by membrane EVs.

Studies have shown that EVs released by cancer cells appear to play an important role in tumor progression, diagnosis, and therapeutic potential^36–38^. The role played by a number of TOP-specific substrates in the proliferation of prostate cancer cells is a topic of interest in cancer research. Early stage prostate cancer proliferation is dependent on male sex hormones (androgens) and can be effectively treated with androgen-ablation therapy. Analogs of TOP substrate GnRH-I have been utilized in the treatment of prostate cancer since the early 1980s^39, 40^, for which they are used to suppress the HPG axis during the early androgen dependent phase of prostate cancer. GnRH antagonists are currently FDA-approved frontline treatments of prostate cancer and are widely used in clinical applications^39^.

As a key modulator of the activities of GnRH and its other physiological substrates, extracellular TOP stands as an important subject of study for further understanding of the pathophysiological role of these hormones and monitoring the progression of prostate cancer. In contrast to the soluble form of TOP, which easily diffuses into the circulation and can be diluted by large volumes of circulating blood, the EV-carried TOP may stay in the tumor microenvironment in a relatively high concentration. Thus, this EV-associated format of TOP may enable the enzyme to work more potently on its substrates, therefore contributing to the progression of pathological conditions. Our study paved the way for further understanding of the activities and functions of EV-associated TOP in the extracellular space and the specific mechanisms by which TOP becomes bound to the plasma membrane and is subsequently released with EVs. Our novel finding may provide insight into the development of more effective therapeutic strategies and monitoring methods in the treatment of prostate cancer.

